# Pitfalls and Remedies for Cross Validation with Multi-trait Genomic Prediction Methods

**DOI:** 10.1101/595397

**Authors:** Daniel Runcie, Hao Cheng

## Abstract

Incorporating measurements on correlated traits into genomic prediction models can increase prediction accuracy and selection gain. However, multi-trait genomic prediction models are complex and prone to overfitting which may result in a loss of prediction accuracy relative to single-trait genomic prediction. Cross-validation is considered the gold standard method for selecting and tuning models for genomic prediction in both plant and animal breeding. When used appropriately, cross-validation gives an accurate estimate of the prediction accuracy of a genomic prediction model, and can effectively choose among disparate models based on their expected performance in real data. However, we show that a naive cross-validation strategy applied to the multi-trait prediction problem can be severely biased and lead to sub-optimal choices between single and multi-trait models when secondary traits are used to aid in the prediction of focal traits and these secondary traits are measured on the individuals to be tested. We use simulations to demonstrate the extent of the problem and propose three partial solutions: 1) a parametric solution from selection index theory, 2) a semi-parametric method for correcting the cross-validation estimates of prediction accuracy, and 3) a fully non-parametric method which we call CV2*: validating model predictions against focal trait measurements from genetically related individuals. The current excitement over high-throughput phenotyping suggests that more comprehensive phenotype measurements will be useful for accelerating breeding programs. Using an appropriate cross-validation strategy should more reliably determine if and when combining information across multiple traits is useful.

## INTRODUCTION

Genomic Selection (GS) aims to increase the speed and accuracy of selection in breeding programs by predicting the genetic worth of candidate individuals or lines earlier in the selection process, or for individuals that cannot be directly phenotyped (Meuwissen *et al*. 2001; Hayes *et al*. 2009; Crossa *et al*. 2017). Genomic selection works by training statistical or Machine Learning models on a set of completely phenotyped and genotyped individuals, and then using the trained model to predict the genetic worth of unmeasured individuals. If the predictions are reasonably accurate, selection intensity can be increased either because the population size of candidate individuals is larger or their true genetic worth is estimated more accurately.

Predictions of genetic values are usually based only on the genotypes or pedigrees of the new individuals. However predictions can in some cases be improved by including measurements of “secondary” traits that may not be of direct interest but are easier or faster to measure (Thompson and Meyer 1986; Pszczola *et al*. 2013; Lado *et al*. 2018). This is one goal of multi-trait genomic prediction. Multi-trait prediction is most useful for increasing the accuracy of selection on a single focal trait when that trait has low heritability, the “secondary” traits have high heritability, and the genetic and non-genetic correlations between the traits are large and opposing (Thompson and Meyer 1986; Jia and Jannink 2012; Cheng *et al*. 2018). With the advent of cheap high-throughput phenotyping, there is great interest in using measurements of early-life or easily accessible traits to improve prediction of later-life or more expensive traits, and multi-trait prediction models are attractive methods for leveraging this information (Pszczola *et al*. 2013; Rutkoski *et al*. 2016; Fernandes *et al*. 2017; Lado *et al*. 2018).

A large number of genomic prediction methods are available, and the best model varies across systems and traits (Heslot *et al*. 2012; de Los Campos *et al*. 2013). Due to their complexity and often high-dimensional nature, genomic prediction methods are prone to overfitting and require regularization to perform well on new data. Therefore, comparing models based on their ability to fit existing data (ex. with *R*^2^) is unreliable; every candidate model could explain 100% of the variation in a typical-size dataset.

Instead, prediction models are generally compared by cross-validation (Meuwissen *et al*. 2001; Utz *et al*. 2000; Gianola and Schon 2016). The basic idea of cross-validation is to separate the model fitting and tuning process from the model evaluation process by using separate datasets for each (Hastie *et al*. 2009). This penalizes models that fit too closely to one data set at the expense of generalization. In this way, cross-validation is meant to accurately simulate the real-world usage of the model: predicting the genetic values of un-phenotyped individuals; i.e. those not available during the model fitting process itself. Rather than requiring new data *per se*, cross-validation works by splitting an existing dataset into non-overlapping “training” and “testing” partitions, fitting the candidate model to the former, and then evaluating it on its accuracy at predicting the latter. Common measures of accuracy include Pearson’s *ρ* or the square root of the average squared error (RMSE) (Daetwyler *et al*. 2013). This process of splitting, training, and predicting is typically repeated several times on the same dataset to get a combined or averaged measure of accuracy across different random partitions of the data.

Estimates of model accuracy by cross-validation are not perfect (Hastie *et al*. 2009). They are subject to sampling error as are any other statistic. They are also typically downwardly biased because smaller training datasets are used for the cross-validation than in the actually application of a model. However in typical cases, this downward bias is the same for competing models and thus does not impact model choice (Hothorn *et al*. 2005).

However, cross-validation can give upwardly biased estimates of model accuracy when misused due to various forms of “data-leakage” between the training and testing datasets, leading to overly optimistic estimates of model performance (Kaufman *et al*. 2012). Several potential mistakes in cross-validation experiments are well known:

- **Biased testing data selection**. The individuals in the model testing partitions should have the same distribution of genetic (and environmental) relatedness to the training population as individuals in the remaining target population (Amer and Banos 2010; Daetwyler *et al*. 2013). For example, if siblings or clones are present in the data, they should not be split between testing and training partitions unless siblings or clones of individuals in the training partition are also at the same frequency in the target population. Similarly, if the goal is to predict into a diverse breeding population, the cross-validation should not be performed only within one F2 mapping population.
- **Overlap between the testing and training datasets**. The observations used as testing data should be kept separate from the training data at all stages of the cross-validation procedure. For example, if data from individuals in the testing dataset are used to calculate estimated genetic values (EBVs) for model training, then the testing and training datasets are overlapping, even if the testing individuals themselves are excluded from model training (Amer and Banos 2010).
- **Pre-selection of features (e.g. markers) based on the full dataset before cross-validation**. All aspects of model specification and training that rely on the observed phenotypes should be performed only on the training partitions, without respect to the testing partition. For example, if a large number of candidate markers are available but only a portion will be included in the final model, the selection of markers (i.e. features) should be done using only the training partition of phenotypes and the selection itself should be repeated each replicate of the cross-validation on each new training dataset. If the feature selection is only done once on the whole dataset before cross-validation begins, this can lead to biased estimates of model accuracy (Hastie *et al*. 2009).

If these mistakes are avoided, cross-validation generally works well for comparing among single-trait methods, and in some cases for multi-trait methods. However, our goal in this paper is to highlight a challenge with using cross-validation to choose between single-trait methods and multi-trait methods; specifically multi-trait methods that use information from “secondary” traits *measured on the target individuals* to inform the prediction of their focal trait(s). In this case, standard cross-validation approaches lead to biased results. As we discuss below, the source of bias is not data leakage between the training and testing data *per se*, but correlated errors with respect to the true genetic merit between the secondary traits in the training data and the focal train in the testing data. Note that this issue only occurs when the multiple traits are measured on the same individuals, and the traits share non-genetic covariance. When traits are measured on different individuals, the standard cross-validation approach is appropriate.

In the following sections, we first describe the opportunity offered by multi-trait genomic prediction models in this setting, and the challenge in evaluating them. We then develop a simulation study that highlights the extent of the problem. Next, we propose three partial solutions that lead to fairly consistent model selections between single and multi-trait models under certain situations. Finally, we draw conclusions on when this issue is likely to arise and when it can be safely ignored.

## GENERAL SETTING

Multi-trait genomic prediction is useful in two general settings: 1) When the overall value of an individual depends on each trait simultaneously (ex. fruit number and fruit size) and these traits are correlated, and 2) When a focal trait is difficult or expensive to measure on every individual, but other correlated traits are more readily available (Thompson and Meyer 1986; Pszczola *et al*. 2013; Lado *et al*. 2018). While multi-trait models are clearly necessary in the first setting, in the second the value of the secondary traits depends on several factors including i) the repeatability of the focal and secondary traits, ii) the correlations among the traits and the *cause* of the correlations (i.e. genetic vs non-genetic), and iii) the relative expenses of collecting data on each trait.

Here we focus on the goal of predicting a single focal trait using information from both genetic markers (or pedigrees) and phenotypic information on other traits. Even within this context, there are also two distinct prediction settings: 1) Predicting the focal trait value for new individuals that are yet to be phenotyped for any of the traits, and 2) Predicting the focal trait value for individuals that have been partially phenotyped; phenotypic values for the secondary traits are known and we wish to predict the individual’s genetic value for the focal trait. These settings were described by (Burgueño *et al*. 2012) as CV1 and CV2, respectively, although those authors focused on multi-environment trials rather than single experiments with multiple traits per individual. The same naming scheme has since been extended to the more general multiple-trait prediction scenarios (Lado *et al*. 2018).

The key difference between CV1 and CV2-style multi-trait prediction is that in the former, the secondary traits help refine estimates of the genetic values of *relatives* of the individuals we wish to predict, while in the latter, the secondary traits provide information directly about the genetics of the target individuals themselves. This direct information on the target individuals is generally useful (as we demonstrate below). However, it comes with a cost for the *evaluation* of prediction accuracy by cross-validation. Since we do not know the true genetic values for the testing individuals, we must either use a model to estimate the genetic values or simply use their phenotypic value as a proxy. Unfortunately, if we use our genetic model to estimate these values, we are breaking the independence between the testing and training data, and therefore have biased estimates of cross-validation accuracy. On the other hand, if we simply use the phenotypic values of the focal trait as our predictand, these may be biased towards or away from the true genetic values depending on the non-genetic correlation between the focal and secondary traits. This leads to either over- *or under*-estimation of the prediction accuracy of our multi-trait models. In realistic scenarios, this can lead users to select worse models.

## MATERIALS AND METHODS

We used a simulation study to explore conditions when naive cross-validation experiments as described above lead to sub-optimal choices between single and multi-trait genomic prediction methods. Our simulations were designed to mimic the process of using cross-validation to compare single and multi-trait models based on their prediction accuracies. We repeated this simulation across scenarios with different genetic architectures for two traits: a single “focal” trait and a single “secondary” trait. Specifically, we modified the heritability and correlation structure of the two traits. These are the most important parameters for determining the relative efficiencies of single- and multi-trait prediction models (Thompson and Meyer 1986). Sample size and level of genomic relatedness will also affect the comparisons, but are likely to only quantitatively (but not qualitatively) change the relative performances of the models and the accuracy of cross-validation.

To make our simulations realistic, we based them on genomic marker data from 803 lines from a real wheat breeding program (Lopez-Cruz *et al*. 2015). We downloaded the genomic relationship matrix **K** based on 14,217 GBS markers from this population. We used this relationship matrix to generate a set of simulated datasets covering all combinations of the following parameters: the relative proportions of genetic and non-genetic variation for each trait *(h*^2^ = {0.2, 0.6}), and the genetic and non-genetic correlations between the traits *ρ*_*g*_ = {0,0.3,0.6}, *ρ*_*R*_ = {−0.6, −0.4, –0.2,0,0.2,0.4,0.6}, drawing trait values for each simulation from multivariate normal distributions. In particular, we set:

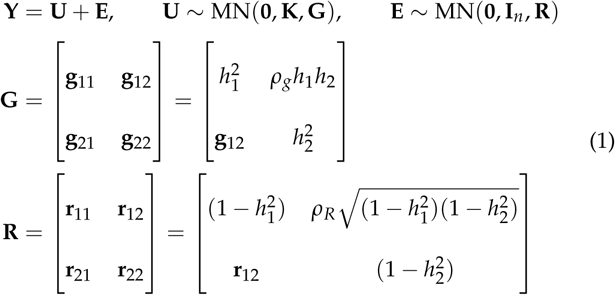

where MN(.) is the Matrix normal distribution, **Y** = [**y**_1_,**y**_2_] are the phenotypic values for the two traits in the *n* individuals, **U** = [**u**_1_, **u**_2_] are the true genetic values for the two traits, and **E** = [**e**_1_, **e**_2_] are the true non-genetic deviations for the two traits. We repeated this process 500 times for each of the 42 combinations of the genetic architecture parameters. To improve the consistency of the simulations, we used the same draws from a standard-normal distribution for all 42 parameter combinations, but new draws for each of the 500 simulations.

After creating the 803 simulated individuals, we randomly divided them into a training partition and a testing partition. We arranged the rows of **Y** so that the testing individuals were first, and correspondingly partitioned **K** into:

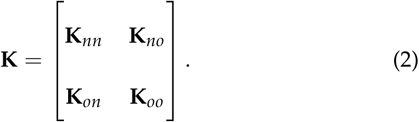

Here and below, the subscript _*n*_ refers to the testing partition (i.e. “new” individuals) and the subscript _*o*_ refers to the training partition (i.e. “old” individuals). We use the hat symbol (^) to denote parameter estimates or predictions.

We then fit single- and multi-trait linear mixed models to the training data and used these model fits to predict the genetic values for the focal trait (trait 1) in the testing partition.

Specifically, for the single-trait method we fit a univariate linear mixed model to the training data **y**_*o*1_:

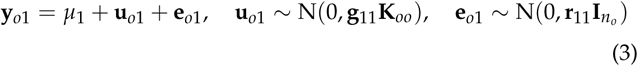

by Restricted Maximum Likelihood using the `relmatlmer` function of R package (Ziyatdinov *et al*. 2018) and extracted the BLUPs **û**_*o*1_. Note: an expanded version of these derivations are provided in the Appendix. We then calculated predicted genetic values for the testing partition **u**_*n*1_ as:

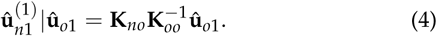

For the multi-trait model, we stacked the vectors of the two traits in the training dataset into the vector 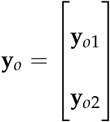 and fit:

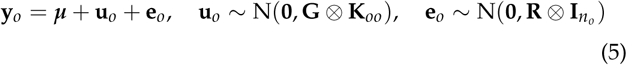

using the `relmatLmer` function, extracted estimates 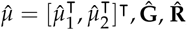, and BLUPs **û**_*o*_.

To make predictions of the genetic values for the focal trait in the testing partition in the CV1 case without use of **y**_*n*2_, we calculated:

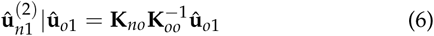

which has the same form as for the single trait model, but the input BLUPs **û**_*o*1_ are different.

To make predictions of the genetic values for the focal trait in the testing partition in the CV2 case, using the phenotypic observations of the secondary trait **y**_*n*2_, we used a two step method. First, we estimated **û**_*o*_ above based on both traits in the training data. Then we combined these estimates with the observed phenotypes of the testing data to calculate genetic predictions for the testing data:

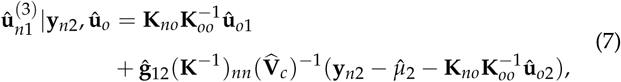

where 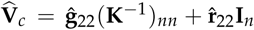. This two-step method will be slightly less accurate than a one-step method that used **y**_*n*2_ during the estimation of **û**_*o*_, but is much easier to implement in breeding programs because no genotype or phenotype data of the evaluation individuals is needed during the model training stage.

We measured the accuracy of these three predictions by calculating the correlation between the prediction 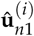 and three predictands over the 500 simulations:

- **u**_*n*1_: The true genetic value.
- **y**_*n*1_: The phenotypic values of the testing individuals.
- **ũ**_*n*1_: The estimated genetic values of the validation individuals using the full dataset (including **y**_*n*1_).

For the second accuracy measure that uses phenotypic values as predictands, we “corrected” the correlations by dividing by the true value of 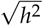 to account for the larger variance of **y**_*n*1_ relative to **u**_*n*1_. This impacts the denominator of the correlation (Daetwyler *et al*. 2013), but since it is the same across methods, does not impact their comparison.

As described below, we also simulated phenotypes for an additional set of individuals **y**_*x*_ not included in either the validation or testing partitions. These individuals were selected to be close relatives of each of the validation partition individuals but experienced different micro-environments.

For each combination of genetic parameters, we declared the “best” prediction method to be the one with the highest average correlation with the true genetic values across the 500 simulations. Then we counted the proportion of the simulations in which this “best” method actually had the highest estimated accuracy when scored against **y**_*n*1._

## Data availability

Scripts for running all simulations and analyses described here are available at https://github.com/deruncie/multiTrait_crossValidation_scripts.

## RESULTS

Although we ran simulations for two levels of heritability for the focal trait 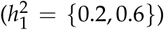 we present results only for 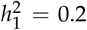. This is the “most-difficult” setting for prediction-when the heritability of the trait is low-but also the setting when we would expect the greatest benefit of using multi-trait models. Results for 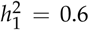 were qualitatively similar, but with higher overall prediction accuracies of all methods.

### Accuracy of single and multi-trait methods in simulated data

With 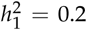 the true accuracy of prediction was moderate for all methods (*cor*(**û**_*n*1_, **u**_*n*1_) ∼ 0.4 − 0.6, Figure 1). Prediction accuracies for the single-trait method were constant across settings with different correlation structures because information from the secondary trait was not used.

**Figure 1.**
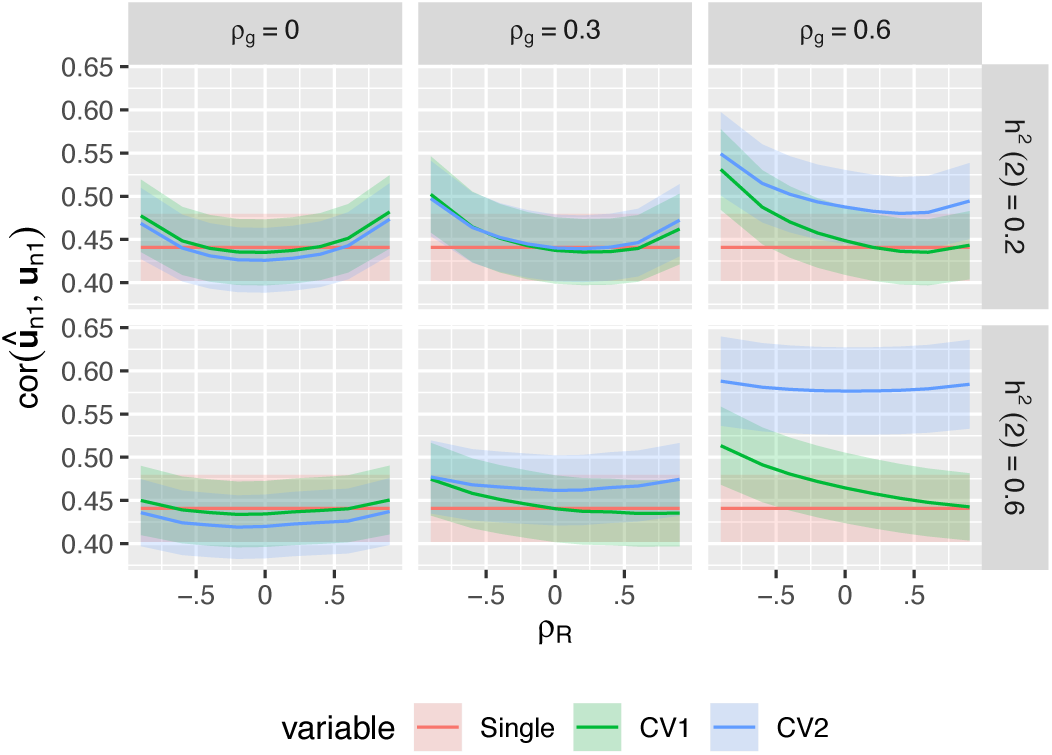
True prediction accuracy of single-trait and multi-trait prediction methods in simulated data. 500 simulations were run for each heritability of the secondary trait 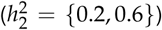, and each combination of genetic and non-genetic correlation between the two traits *(ρ*_*g*_*=* {0,0.3,0.6},*ρ*_*R*_ = {−0.6, −0.4, −0.2,0,0.2,0,4,0.6}), all with 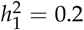. For each simulation, we used 90% of the individuals as training to fit linear mixed models (either single or multi-trait), predicted the genetic values of the remaining validation individuals, and then measured the Pearson’s correlation between the predictec (**û**_*n*1_) and true (**u**_*n*1_) genetic values. In the CV1 method, we used only information on the training individuals to calculate **û**_*n*1_-In the CV2 method, we used the training individuals to calculate **û**_*o*_ and combined this with the observed phenotypes for the secondary trait on the validation individuals (**y**_*n*2_). Curves show the average correlation for each method across the 500 simulations. Ribbons show ±1.96 × *SE* over the 500 simulations.

The “standard” muti-trait model (i.e. CV1-style) that used phenotypic information only on the training partition slightly out-performed the single-trait model in some settings, more-so when the genetic and non-genetic correlations between traits were large and opposing and when the genetic determinacy of the secondary trait was high (Thompson and Meyer 1986). However it performed slightly worse whenever the genetic and residual correlations between traits were low. This was caused by inaccuracy in the estimation of the two covariance parameters 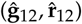. Neither multi-trait model performed worse than the single-trait model when the true **G** and **R** matrices were used Supplemental Figure 1, which we also verified by calculating the expected prediction accuracies analytically (See Appendix). In real data, multi-trait models require estimating more (co)variance parameters and therefore can show reduced performance when data are limited.

The CV2-style multi-trait method, which leverages additional phenotypic information on the secondary trait from the testing partition itself, showed dramatic improvements in prediction accuracy whenever genetic correlations among traits were large, irregardless of the non-genetic correlation between the traits. This is similar to the benefits seen by (Rutkoski *et al*. 2016) and (Lado *et al*. 2018). When the heritability of the secondary trait was high, the improvement in prediction accuracy was particularly dramatic (increasing to ∼ *ρ* = 0.6). This is the potential advantage of incorporating secondary traits into prediction methods. However, the CV2 method also requires estimating **G** and **R**, and its performance was lower than the single-trait method whenever both genetic and residual correlations were low.

Therefore, multi-trait methods will not always be useful and it is important to test the relative performance of the different methods in real breeding scenarios. Unfortunately, we never know the true genetic values (**u**_*n*1_), and so must use proxy predictands to evaluate our methods in real data (Daetwyler *et al*. 2013; Legarra and Reverter 2018). In Figures 2A-B, we compare the prediction accuracies of the three methods using two candidate predictands: the observed phenotypic values (**y**_*n*1_) and estimated genetic values from a joint model fit to the complete dataset **(ũ**_*n*1_).

Using the observed phenotypic values (**y**_*n*1_) as the predictand, the estimated accuracy of both the single-trait and CV1-style multi-trait prediction methods consistently under-estimated their true prediction accuracies. This is expected because in this setting 80% of the phenotypic variation is non-genetic and cannot be predicted based on relatives alone. We therefore follow common practice to report a “corrected” estimate of the prediction accuracy by dividing by 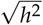 in Figure 2A. This correction factor itself must be estimated in real data, but when comparing models the same value of *ĥ*^2^ should be used for each model so that differences in these estimates do not bias model selection.

**Figure 2.**
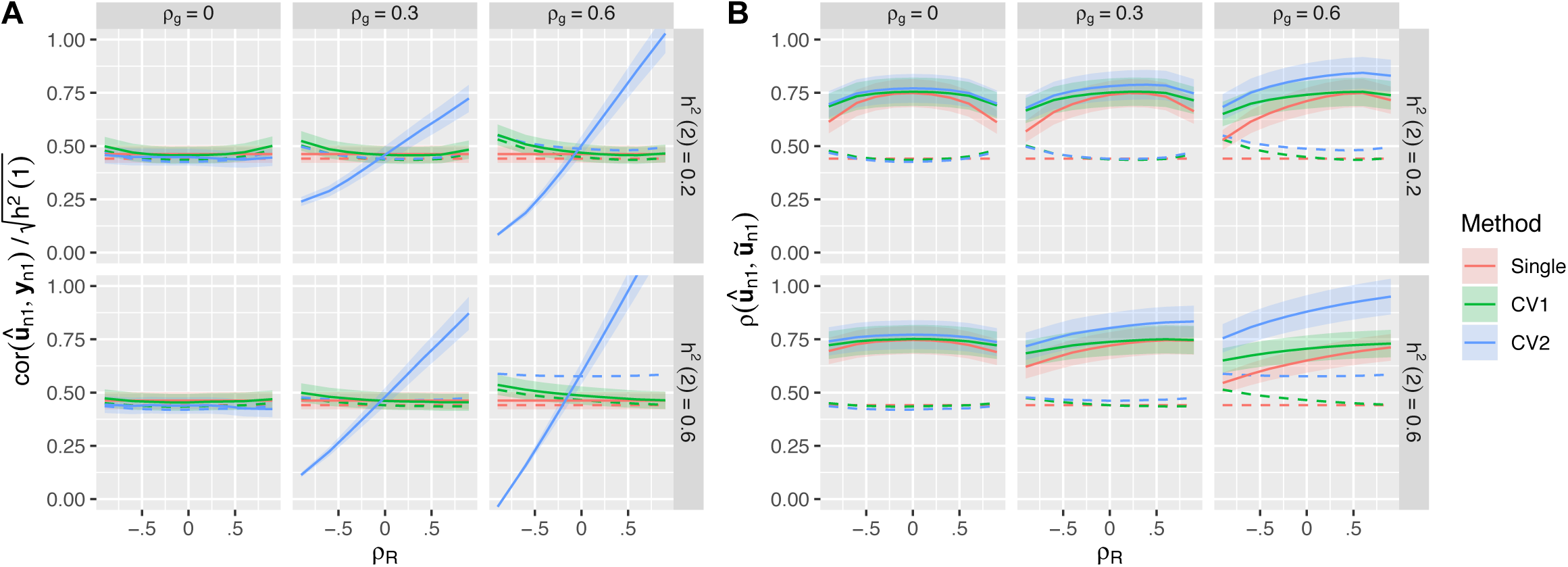
Estimated prediction accuracies based on candidate predictands. For the same set of simulations described in Figure 1, we estimated the prediction accuracies of the three methods using two different candidate predictands: **(A)** The observed phenotypic value **y**_*n*1_ for each training individual (with the correlation corrected by 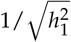), or **(B)** An estimate of the genetic value of each training individual based on BLUPs calculated using the complete phenotype data (**ũ**_*n*1_). Solid lines in each panel show the average *estimated* accuracy for each method across the 500 simulations. Ribbons show ±1.96 × *SE* over the 500 simulations. Dotted lines show the average *true* accuracy from Figure 1.

In contrast, the estimated accuracy of the CV2-style multi-trait method varied dramatically across simulated datasets. We tended to overestimate the true accuracy when both genetic and non-genetic correlations were large and in the same direction, and dramatically underestimate the true accuracy when the two correlations were opposing. Importantly, there are situations where the CV2-style method appears to perform worse than the single-trait method based on **y**_*n*1_ but actually performs better. Therefore, the observed phenotypic values are not reliable predictands to evaluate CV2-style methods when the intent is to estimate true genetic values and *ρ*_*R*_ ≠ 0.

On the other hand, using estimated genetic values from a joint model fit to the complete dataset (**ũ**_*n*1_) as the predictand led to dramatic over-estimation of the true prediction accuracy for all methods. This is also expected because the training data are used both to train the prediction model *and also* to create the testing dataset, a clear violation of the cross-validation rules that these datasets must be kept separate at all stages of the analysis. Again, the bias was most severe for the CV2-style method. Since this method is clearly invalid, we do not consider it further.

### Effects of predictand on model selection

To demonstrate the impact of biased estimates of model accuracy using **y**_*n*1_ on the effectiveness of model selection, we assessed in each simulation whether the single-trait or multi-trait methods had a higher *estimated* accuracy, and compared this result to the *true* difference in prediction accuracies in that simulation setting.

Figure 3 shows that selecting between the single-trait and CV1-style multi-trait models based on estimated accuracy using **y**_*n*1_ generally works well. Whenever one method is clearly better, we are able to choose that method > 50% of the time. But we never choose correctly < 50% of the time, even when the methods are approximately equivalent.

**Figure 3.**
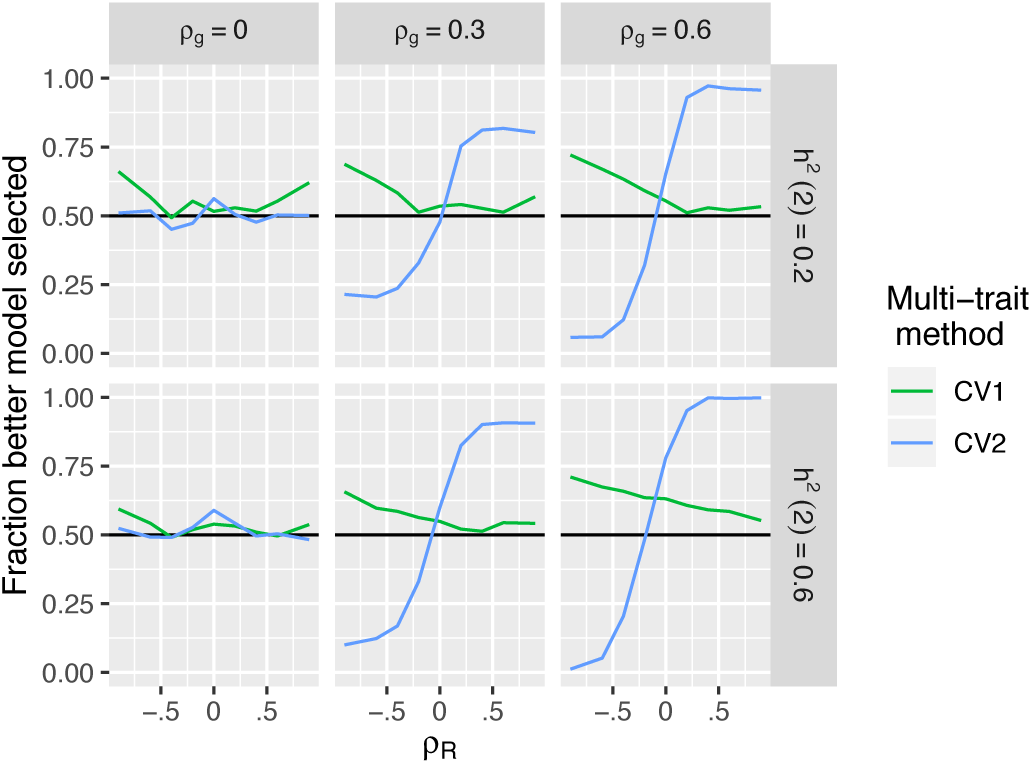
Impact of using phenotypic data to select between single-trait and multi-trait prediction methods. For each of the 500 simulations per genetic architecture described in Figure 1, we compared the estimated accuracy of a multi-trait prediction to the single-trait prediction. We then calculated the fraction of times that the selected model had higher average true accuracy in that setting (as shown in Figure 1).

In contrast, when selecting between the single-trait and CV2-style multi-trait methods based on estimated accuracy using **y**_*n*1_, the differential bias in estimated accuracy between the two methods frequently lead to sub-optimal model selection (Figure 3B). With opposing genetic and non-genetic covariances between the two traits, the better model was chosen < 10% of the time. In these situations, using **y**_*n*1_ to select a prediction method will obscure real opportunities to enhance prediction accuracy using multi-trait prediction models.

### Alternative estimates of multi-trait prediction accuracy

The CV2-style prediction method can be powerful because **y**_*n*2_ provides information on the genetic value of the testing individuals themselves (through **u**_n2_), while **y**_*o*1_ only provides indirect information on the genetic values of the testing individuals through the relatives. However, estimating prediction accuracy using **y**_*n*1_ fails for the CV2-style prediction method because both the focal and secondary traits are observed on the same individual and therefore share the same non-genetic sources of variation. Since the CV2 method uses **y**_*n*2_, non-genetic deviations for the secondary trait **e**_*n*2_ push **û**_*n*1_ either towards or away from **y**_*n*1_ depending on the estimated correlation 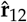. This either inflates or deflates the estimated accuracy, leading to incorrect model choices.

We now compare the effectiveness of three strategies for estimating cross-validation accuracy of CV2-style methods. To our knowledge, the second and third strategies are novel. Because the three methods have different data requirements, we implemented different experimental designs for each evaluation strategy.

### Parametric estimate of accuracy

Our prediction **û**_*n*1_ is similar to a selection index because it combines multiple pieces of information into a linear prediction. The accuracy of an index **I** is: 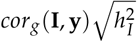, the genetic correlation between the index and phenotype multiplied by the heritability of the index (Falconer and Mackay 1996; Lopez-Cruz *et al*. 2019). Neither the genetic correlation nor the heritability can be directly observed, but we can estimate both as parameters of a multi-trait linear mixed model with the same form as (5). To be a valid cross-validation score, these parameters must be estimated with data only in the validation partition, rather than reusing estimates from model training. Since both model training and model evaluation equally require estimates of **G** and **R**, we divided the data 50:50 into training and validation partitions in each simulation, thus using 404 lines to train the prediction models and 403 lines to evaluate the prediction accuracy.

The parametric estimates of prediction accuracy for the CV2 method were less biased than the 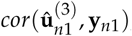, the non-parametric estimates using **y**_*n*1_ as a predictand (Figure 4A, compare to Figure 2). This led to more consistent model selections between the CV2 and single-trait methods (Figure 4B). However, the parametric approach still underestimated the accuracy of the CV2 method when the genetic and residual correlations were in opposite directions, leading to model selection accuracies <50%. This negative bias was due to poor estimation of **G** and **R** for the selection indices, given the limited sample sizes remaining after the data were partitioned.

**Figure 4.**
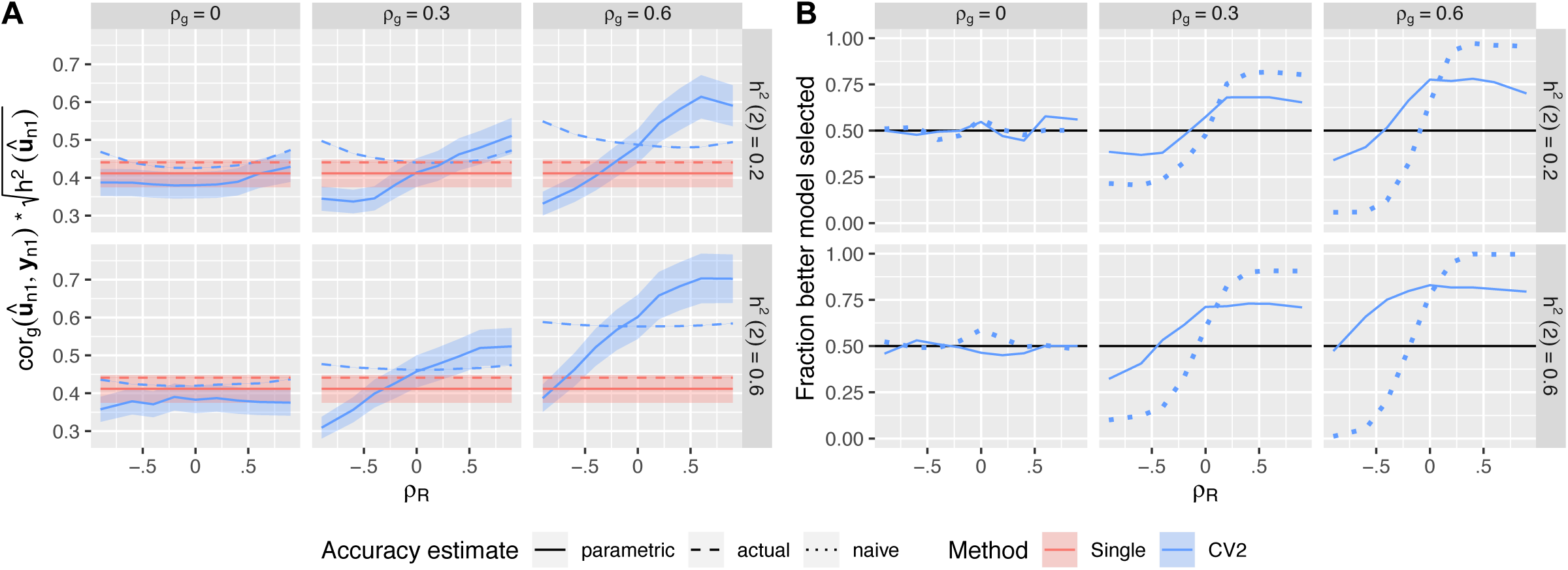
Parametric accuracy estimates. Estimated prediction accuracies and model selection accuracies for CV2-style methods using the parametric method. **(A)** Solid curves: estimates of prediction accuracy. Dashed curves: true prediction accuracy based on **u**_*n*1_ Dotted curves: estimated prediction accuracy using **y**_*n*1_ from Figure 2A. Ribbons show ±1.96 × *SE* over the 500 simulations. **(B)** Solid curves: Fraction of the 500 simulations in which the better method (between CV2 and single-trait) for predicting the true genetic values was correctly selected. Dotted curve: model selection based on the naive prediction accuracy.

### Semi-parametric estimate of accuracy

In principle, we can correct for the bias in the non-parametric accuracy estimate 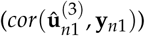 from the CV2-style method by calculating an adjustment factor based on the theoretical bias relative to the true accuracy 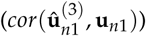 This is similar to the semi-parametric accuracy estimates presented by (Legarra and Reverter 2018), and the “correction” of accuracy estimates by 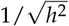 used above to account for the difference in variance between **y**_*n*1_ and **u**_*n*1_. As we derive in the Appendix, the difference between the true correlation from a CV2-style methods and its CV2 cross-validation estimate when a single secondary trait is used is:

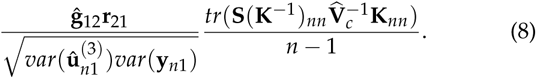

with **V**_*c*_ defined above and 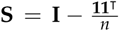. The bias is a function of the the correlation among traits through the product **ĝ**_12_**r**_21_(as the second term does not involve these parameters, and in most cases is ≈ 1), and is large and positive (i.e. accuracy is overestimated) when **ĝ**_12_ and **r**_12_ are large and in the same direction, and large and negative (i.e. accuracy is underestimated) when these covariances are in opposite directions. Given this result, we can correct 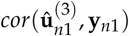 by subtracting 8 from the estimated correlation, again corrected by 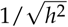 (Figure 5).

**Figure 5.**
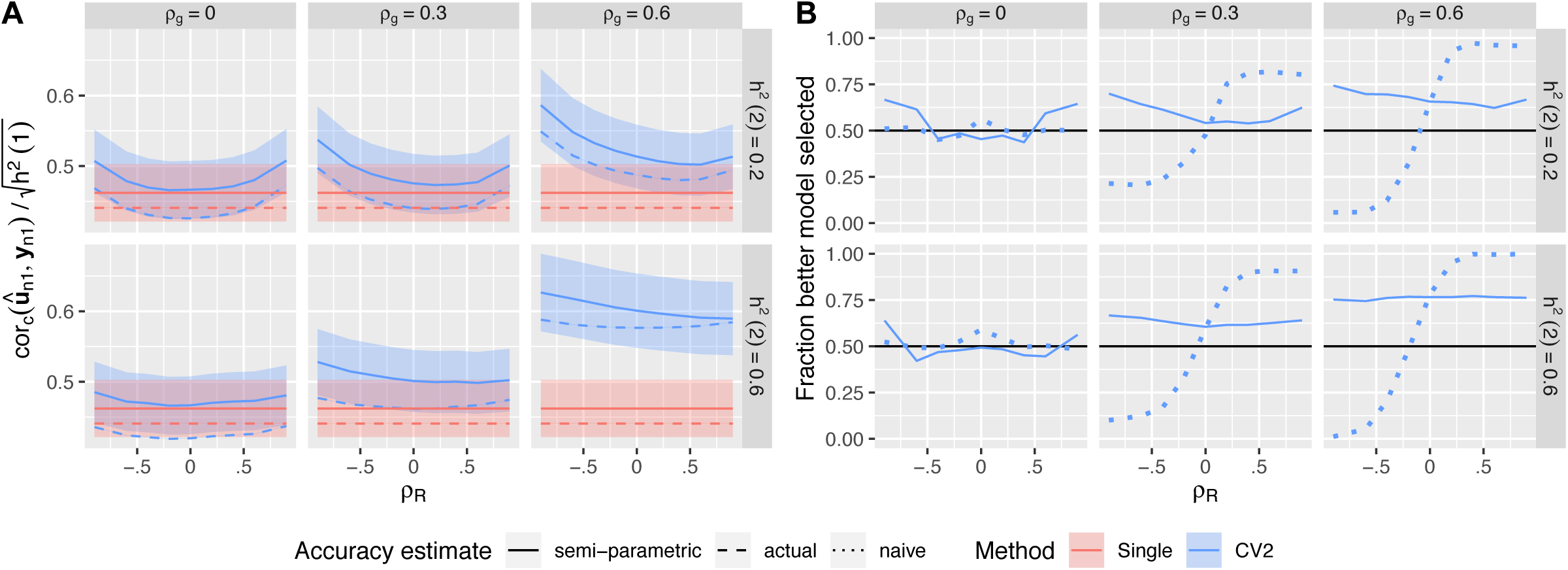
Semi-parametric accuracy estimates. Estimated prediction accuracies and model selection accuracies for CV2-style methods after semi-parametric correction. **(A)** Solid curves: corrected estimates of prediction accuracy. Dashed curves: uncorrected estimates of prediction accuracy based on **y**_*n*1_ (mirroring Figure 3). Dotted curves: true prediction accuracy. Ribbons show ±1.96 × *SE* over the 500 simulations. **(B)** Solid curves: Fraction of the 500 simulations in which the better method (between CV2 and single-trait) for predicting the true genetic values was correctly selected. Dotted curve: model selection based on the naive un-corrected prediction accuracy.

Clearly, the quality of this correction will depend on the accuracy of ĝ_12_ and 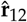 as estimates of **g**_12_ and **r**_12_-In Figure 5A, we show that the corrected correlation estimate has greatly reduced bias, particularly the dependence of the bias on the non-genetic covariance between the traits **r**_12_. However the correction is not perfect. Corrected accuracy estimates tend to overestimate the true accuracy. This over-estimation is caused by error in **Ĝ** and 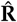. as estimates of the true covariances: The correction factor is nearly perfect when the true covariance matrices are used in place of their estimates Supplemental Figure 2.

Using the semi-parametric accuracy estimates, we are more successful at selecting the best model over the range of genetic architectures (Figure 5B). The frequency of selecting the correct model rarely drops below 50% and is relatively constant with respect to the residual correlation between traits.

### CV2* cross-validation strategy

Since the biased estimate of prediction accuracy for CV2-style methods is due to non-genetic correlations between **y**_*n*2_ used for prediction and the predictand **y**_*n*1_, an alternative strategy, which we call CV2*, is to use phenotypic information on close relatives of the testing individuals (**y**_*x*1_) to validate the model predictions in place of their own focal trait phenotypes (**y**_*n*1_). These “surrogate” validation individuals must also be excluded from the model training and raised so that they do not share the same non-genetic deviations as the testing individuals: *car*(**e**_*x*1_,**e**_*n*1_) = 0. Therefore, **û**_*x*1_ will not be artificially pushed towards or away from **u**_***x*1**_ (measured on relatives) by **y**_*n*2_ (measured on testing individuals), preventing this source of bias in the estimated accuracy.

We implemented the CV2* cross-validation strategy in two ways, simulating two different breeding schemes.

First, we considered the situation common in plant breeding where inbred lines (i.e. clones) are tested, and each line is grown in several plots in a field Bernardo (2002). Here, we can use one set of clones for prediction (**y**_*n*2_), and the other set of clones as trait-1 surrogates (**y**_*x*1_). Since they are clones, **u**_*x*1_ = **u**_*n*1_ and **y**_*y*2_ is just as good for predicting **u**_*x*1_ as **y**_*x*2_. Generally in this type of experiment, replicate plots of each line will be combined prior to analysis into a single line mean (or BLUP). But since we require **y**_*n*2_ and **y**_*x*1_ to be recorded from separate individuals, each value will have 2× the residual variance because it is based on 1/2 as much data as the line means used for model training. Therefore, in our simuulations we drew two independent residual values for each line in the validation partition, each with a variance of **2R**. For these simulations, we used a 90:10 training:validation split.

Second, we considered the situation more common in animal breeding where clones are not available. In this case, the best option for CV2* would be to select pairs of closely related individuals to include in the training set; we use the first individual of the pair as **y**_*n*2_ and the second as **y**_*x*1_ To implement this strategy, we again started with a validation partition of 10% of the lines. Then for each line, we selected the most closely related remaining line (arg max_*j*_ **K**_*ij*_ for validation line *i*) and held this additional set of 10% of the lines as **y**_*x*1_. This left a training partition with only 80% of the lines. The average genetic relatedness of validation partition pairs in these simulations was 0.38.

Figure 6A shows that for the first setting with split clones, estimates of prediction accuracy for CV2-style predictions by CV2* are vastly more accurate than the naive estimates based on **y**_*n*1_, but they are slightly downwardly biased because of the increased residual variance of **y**_*n*1_ and **y**_*x*2_. Model selection works fairly well across all settings when clones are used (Figure 6B, blue lines), although with slightly lower success rates than for the semi-parametric method. However, when we implementing the second approach with nearest relatives (not clones), model selection was rarely successful - we consistently chose the wrong model across most simulation settings unless the genetic and residual correlations were opposing. This is because the validation pairs were too distantly related to provide any additional information on genetic merit relative to individuals in the training partition. Interestingly, this method is relatively successful in the situations where the parametric method fails (see Figure 4B), and so may be complimentary.

**Figure 6.**
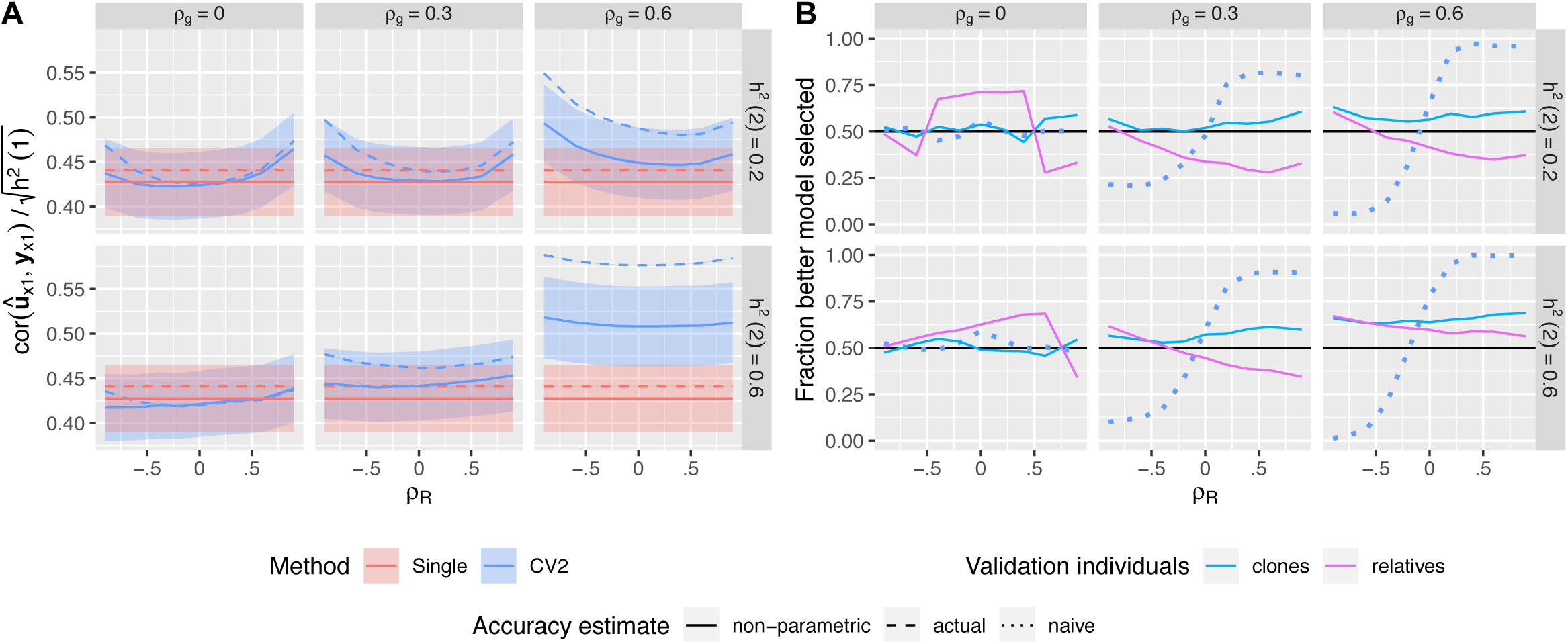
Non-parametric CV* accuracy estimates. Estimated prediction accuracies and model selection accuracies based on the phenotypic values of close relatives. **(A)** Solid curves: Estimated prediction accuracies of the CV2-style and Single-trait methods evaluated against **y**_*x*1_ using clones. Dashed curves: True prediction accuracies of each method. Ribbons show ±1.96 × *SE* over the 500 simulations. **(B)** Solid curves: Fraction of the 500 simulations in which the better method (between CV2 and single-trait) for predicting the true genetic values was correctly selected based on the phenotypes of relatives of the testing individuals. Dotted curve: Fraction of correct models selected based on the naive estimator.

## DISCUSSION

Our study highlights a potential pitfall in using cross-validation to estimate the accuracy of multi-trait genomic prediction methods. When secondary traits are used to aid in the prediction of focal traits and these secondary traits are measured on the individuals to be tested, cross-validation evaluated against phenotypic observations can be severely biased and result in poor model choices. Unfortunately, we rarely know the true genetic value of any individual and therefore can only evaluate our models with phenotypic data (since multi-trait-derived estimated genetic values are even more severely biased as we demonstrated above (Figure 2B)). We cannot find earlier discussions of this problem in the literature. However a growing number of studies aim to use cheap or early-life traits to improve predictions of genetic worth for individuals in later-life traits (ex. Pszczola *et al*. 2013; Rutkoski *et al*. 2016; Fernandes *et al*. 2017; Lado *et al*. 2018). Therefore the issue is becoming more important.

The problematic bias in the cross-validation-based accuracy estimates is caused by non-genetic correlations between the predictors that we want to use (i.e. the secondary traits) and our best predictand (the phenotypic value of the trait in the testing individuals) - non-genetic correlations between two traits measured on the same individual are expected. However, in some cases this correlation is zero by construction, and standard cross-validation approaches can be valid. For example, in the original description of the CV2 cross-validation method by (Burgueño *et al*. 2012), each trait was measured in a different environment. In this case, the traits were measured on different individuals and therefore did not share any non-genetic correlation. Also, CV1-style methods do not suffer from this problem because phenotypic information on the secondary traits *in the testing individuals* is not used for prediction. Similarly, this bias does not occur when the target of prediction is the phenotypic value itself (rather than the individual’s genetic value). For example, in medical genetics the aim is to predict whether or not a person will get a disease or not, not her genetic propensity to get a disease had she been raised in a different environment (ex Spiliopoulou *et al*. 2015; Dahl *et al*. 2016).

We note that the common strategy of two-step genome selection: using single-trait methods to calculate estimated genetic values for each line:trait and then using these estimated genetic values as training (and validation) data, does not get around the problem identified here. Using estimated genetic values instead of phenotypic values will tend to increase the genetic repeatability of the training and validation values, and therefore increase the overall prediction accuracy of all methods. But these estimated genetic values will still be biased by the non-genetic variation, and the biases across traits will still be correlated by the non-genetic correlations. Therefore the same issue will arise.

Also, while we have used a GBLUP-like genomic prediction method for the analyses presented here, the same result will hold for any multi-trait prediction method that aims to use information from **y**_*n*2_ when there are non-genetic correlations with **y**_*n*l_,i.e. any method that is evaluated with the CV2 cross-validation method on multiple traits measured on the same individual (Calus and Veerkamp 2011; Jia and Jannink 2012; Fernandes *et al*. 2017). This includes multi-trait versions of the Bayes Alphabet methods (Calus and Veerkamp 2011; Cheng *et al*. 2018), or neural network or Deep Learning methods (Montesinos-López *et al*. 2018).

We presented three partial solutions to this problem, spanning from fully parametric to fully non-parametric.

The parametric solution relies on fitting a new multi-trait mixed model to the predicted values and the predictand, with the accuracy estimated as the genetic correlation scaled by the heritability of the prediction. This solution is always available as long as the individuals in the validation partition have non-zero genomic relatedness and the full dataset is large enough to estimate genetic correlations in both training and validation partitions. However it generally worked poorly in our simulations because **G** and **R** were not estimated accurately. It may work better with very large datasets. Also, because this parametric approach relies on the same assumptions about the data (i.e. multivariate normality) as the prediction model, it loses some of the guarantees of reliability that completely non-parametric cross-validation methods can claim.

The semi-parametric solution aims to correct the non-parametric correlation estimate for the bias caused by the non-null residual correlation among traits. This correction factor is only needed for CV2-style multi-trait prediction approaches, and is similar to the approach of (Legarra and Reverter 2018) for single-trait models. We show that this correction factor can work well, particularly if the covariances among traits are well estimated. We only derived this correction method for prediction methods based on linear mixed effect models with a single known genetic covariance structure (i.e. GBLUP and RKHS-style methods with fixed kernels), although the approximation 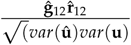 will probably be approximately correct for other methods. However, when covariances are poorly estimated, the correction factor can still lead to biased estimates of model accuracy. We are currently investigating whether Bayesian methods that sample over this uncertainty can be useful, and will implement this method in JWAS (Cheng *et al*. 2018). This method is semi-parametric, so also relies on distributional assumptions about the data and may fail when these assumptions are not met.

As a third alternative, we proposed the CV2* cross-validation method, a fully non-parametric approach for assessing CV2-style multi-trait prediction accuracy. CV2* uses phenotypic values of the focal trait from relatives of the testing individuals in place of the phenotypic values of that trait from the testing individuals themselves. If the close relatives are raised independently, they will not share non-genetic variation, removing the source of bias in the cross-validation estimate (Figure 6A). The CV2* method works best when clones of the testing individuals are available. With clones, secondary trait phenotypes of the testing individuals can be used directly to predict focal trait genetic values of their clones because the genetic values are identical. Replicates of inbred lines are frequently used in plant breeding trials (Bernardo 2002). In this case, all replicates should be held-out as a group from the training data. Then the replicates can be partitioned again into two sets; secondary trait phenotypes from one set can be incorporated into the genetic value predictions for the lines, and these predictions evaluated against the phenotypic values of the other set. To compare this estimate of CV2-style prediction accuracy to the prediction accuracy for a single-trait method, the single-trait method’s predictions should be compared against the same set of replicates of each line (i.e., not a joint average over all replicates of the line as would be typical for single-trait cross-validation). However, because of the separation of the replicates, each replicate will have higher residual variance, which reduces the accuracy of this method. Clones are less common outside of plant breeding, so more distant relatives need to be used instead. In this case, the estimated prediction accuracies of CV2-style methods will be downwardly biased. In our simulations, despite relatively close relatives for each validation line being available, this approach was not successful.

In our simulations, the semi-parametric approach was the most reliable, and the fully parametric approach the least reliable. However the fully parametric approach is always possible to implement while our semi-parametric and non-parametric approaches may not be possible depending on the prediction model used and the structure of the experimental design.

## CONCLUSIONS

We expect that multi-trait methods for genomic prediction carry great promise to accelerate both plant and animal breeding. However there is a need to design better methods to evaluate and train the prediction methods to ensure that models can be accurately compared. We have presented and compared three contrasting methods to evaluating multi-trait methods. Each of these methods is preferred to naive cross-validation when secondary traits of the target individuals are used to predict their focal traits. However the methods can give contrasting answers for different datasets, so careful consideration of which evaluation method to use is critical when choosing among prediction methods.

## ACKNOWLEDGMENTS

We would like to thank Erin Calfee and Graham Coop for suggesting the CV2* method, Gustavo de los Campos for pointing us towards the parametric approach, and helpful comments from two anonymous reviewers.

HC’s work is support by US Department of Agriculture, Agriculture and Food Research Initiative National Institute of Food and Agriculture Competitive Grant No. 2018-67015-27957 DER was supported by the United States Department of Agriculture (USDA) National Institute of Food and Agriculture (NIFA), Hatch project 1010469.

## SUPPLEMENTAL FIGURES

**Supplemental Figure 1 Actual prediction accuracy of single-trait and multi-trait prediction methods in simulated data when G and Rare known**. 500 simulations were run for each heritability of the secondary trait 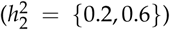, and each combination of genetic and non-genetic correlation between the two traits (*ρ*_*g*_ = {0,0.3,0.6},*ρ*_*R*_ = {−0.6,−0.4,−0.2,0,0.2,0,4,0.6}), all with 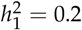. For each simulation, we used the 900 training individuals to fit linear mixed models (either single or multi-trait) conditioning on the true values for **G** and **R**, predicted the genetic values of the 100 testing individuals, and then measured the Pearson’s correlation between the predicted (**û**_*n*1_) and true (**u**_*n*1_) genetic values. In the CV1 method, we used only information on the testing individuals to calculate **û**_*n*1_. In the CV2 method, we used the training individuals to calculate **û** _***o***_ and combined this with the observed phenotypes for the secondary trait on the testing individuals (**y**_*n*2_). Curves show the average correlation for each method across the 500 simulations. Ribbons show ±1.96 × *SE* over the 500 simulations. Dashed lines show analytical calculations of the expected correlation given one representative training:validation data partition.

**Supplemental Figure 2 Estimated prediction accuracies and model selection accuracies for single-trait and multi-trait prediction methods after semi-parametric correction when G and Rare known**. Ribbons show ±1.96 × *SE* over the 500 simulations. Dashed lines show the mean actual prediction accuracy: *cor*(**û**_*n*1_, **u**_*n*1_).

## Appendix

Here, we derive the genomic predictions **û**_*n*1_ given **y** for the three prediction models that we use in the main text, and then evaluate the expected covariances between these predictions and the predictands **u**_*n*1_ and **y**_*n*1_. We derive these relations for the more general situation with *p* ≥ 1 “secondary” traits and a single “focal” trait.

We start with a phenotypic data matrix **Y** with *n* individuals and *p* +1 traits, where the first trait (first column of **Y**) is the “focal” trait, and the other *p* traits are “secondary” traits. We first divide **Y** into a training partition (“old” individuals) and a testing partition (“new” individuals), and arrange them with the testing partition first, so we can partition 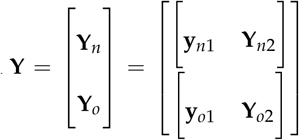. We then work with stacked versions of these phenotype matrices: **y** = *vec*(**Y**),**y**_*n*_ = *vec*(**Y**_*n*_),**y**_*o*_ = *vec*(**Y**_*o*_). Our genetic model for **y** is:

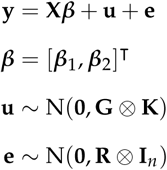

where **G** and **R** are genetic and phenotypic covariance matrices for the *p* +1 traits, and **K** is the *n* × *n* genomic relationship matrix among the lines. For convenience below, we partition the following matrices as follows: We partition the trait vectors for the training individuals and covariance matrices between the “focal” (index 1) and “secondary traits” (index 2):

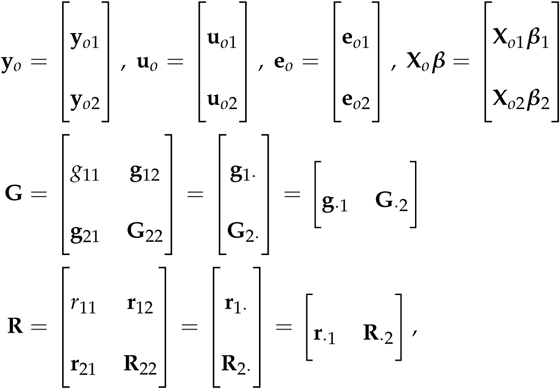

where scalars are normal text, vectors are bold-face lower case letters, and matrices are bold-face capital letters. Partitions for the testing individuals are similar. We also partition the genomic relationship matrix and its inverse between the training and testing individuals:

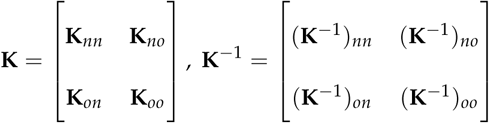

### Derivation of genomic predictions

#### Single trait predictions

For the single-trait prediction, we begin by estimating 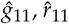 and 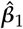 by REML using only **y**_*o*1_. The joint distribution of **u**_*n*1_ and **y**_*o*1_ is:

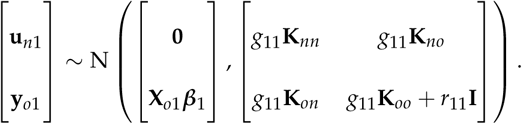

Let: **V**_*o*1_ = *g*_11_**K**_*oo*_ + r_11_**I**. Therefore 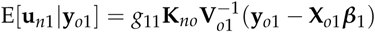, so our prediction is:

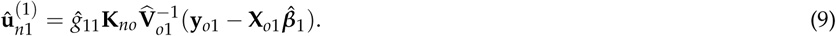

To simplify, note that the joint distribution of **u**_***o*1**_ and **y**_*o*1_ in the training data is:

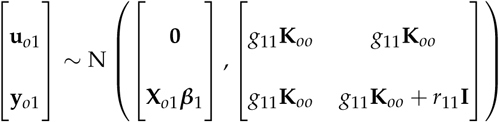

Therefore, 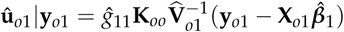. Rearranging and plugging this in above simplifies to: 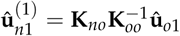.

#### CV1-style multi-trait predictions

For CV1-style multi-trait prediction, we begin by estimating 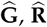. and 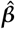 by REML using **y**_*O*_. The joint distribution of **u**_*n*1_ and **y**_*o*_ is:

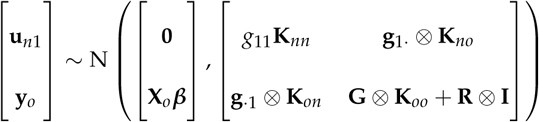

Let **V**_***o***_ = **G⊗ K**_***oo***_ + **R** ⊗ **I**. Therefore, 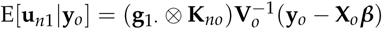, so our prediction is:

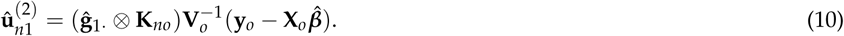

As above, to simplify this expression, we form the joint distribution of **u**_*o*_ and **y**_*o*_ in the training data as:

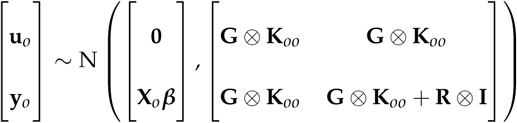

Therefore,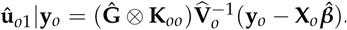. Rearranging and plugging this in above simplifies to: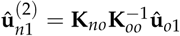.

#### CV2-style multi-trait predictions

For our CV2-style multi-trait prediction, we take a two-step approach. We first estimate **û**_*o*_ from the training individuals and then supplement this with **y**_*n*2_ from the testing individuals. The joint distribution of **u**_*n*1_, **y**_*n*2_ and **u**_*o*_ is:

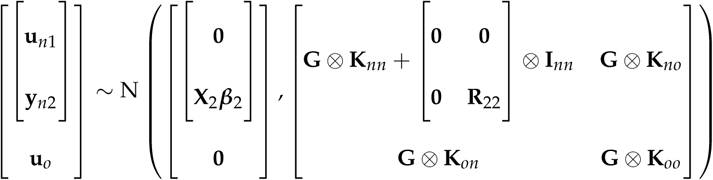

Conditional on a known value of **u**_*o*_ from the training individuals, the distribution of 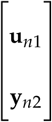 would be:

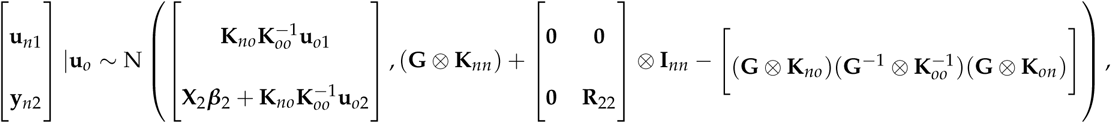

which simplifies to:

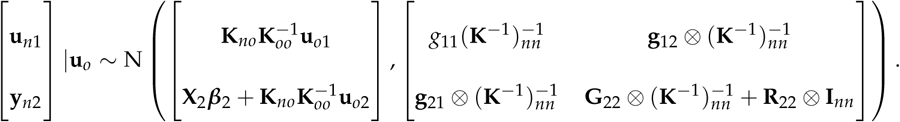

Let 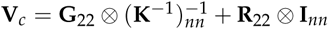. Now, conditioning on observed values of both **u**_*o*_ from the training data and **y**_*n*2_ from the testing data, the expectation of **u**_*n*1_ would be:

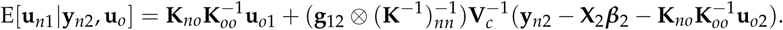

Using this, we form our prediction as:

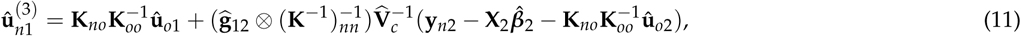

where **û**_*o*1_ and **û**_*o*2_ are extracted from the calculation of **û**_*o*_ for the CV1-style prediction. Plugging in the solutions for these values expands to:

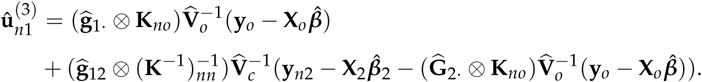

### Expectations of prediction accuracy

Now, we evaluate the expected correlation between a random sample of pairs of elements from our three candidate predictions and the predictand **y**_*n*1_. We compare these expected correlations with the expected “true” correlations with **u**_*n*l_. Below, let *var*(*x*) denote the variance of a random sample from a random vector **x;** *cov*(**x**, **y)** and *cor*(**x**, **y)** denote the covariance and correlation between a random sample of pairs of elements from **x** and **y;** and Cov(**x**, **y)** denote the covariance matrix between vectors **x** and **y**. We use the following results:

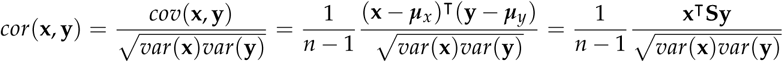

where 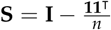.

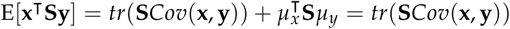

where *tr*(·) is the matrix trace, and ***µ***_*x*_ = **0** and/or ***µ***_*y*_ = **0**. Therefore, the expected correlation between **x** and **y** is approximately:

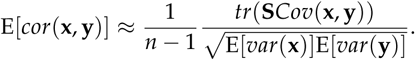

Our goal with cross-validation is to estimate cor(**û**_*n***1**_, **u**_*n*1_). Since we do not know **u**_*n*1_, we approximate the correlation with *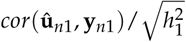*. The factor of 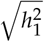 corrects the correlation for the larger variance of **y**_*n*l_ relative to **u**_*n*1_. Otherwise, any difference between these two correlations must be due to their numerators: *tr*(**S***Cov*(**û**_*n*1_, **u**_*n*1_)) and *tr*(**S***Cov*(**û**_*n*1_, **y**_*n*1_)). Thus, for each of the three prediction methods we compare these two numerators to evaluate the accuracy and bias in the approximation.

### Single trait predictions

The numerator of the expected correlation between 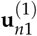 andthetrue genetic values **u**_*n*1_ is:

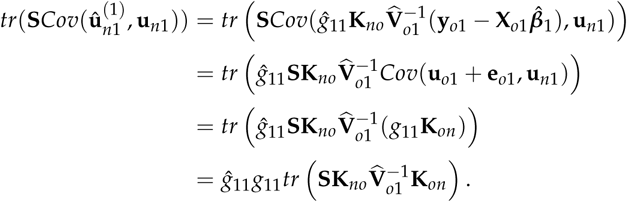

where we assume that 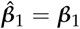, and *Cov*(**e**_*o*1_, **u**_*n*1_) = **0**. The same result for the numerator of the expected correlation between 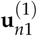 and the observed phenotypic values **y**_*n*1_ is:

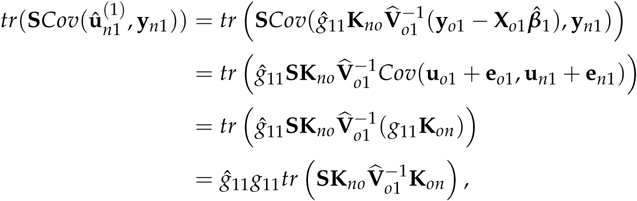

where we additionally assume *Cov*(**u**_*o*1_,**e**_*n*1_) = **0** and *Cov*(**e**_*o*1_, **e**_*n*1_) = **0**. Therefore, the numerators are the same, and 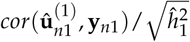 is a consistent estimator for 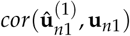.

### CV1-sty/e multi-trait predictions

The numerator of the expected correlation between 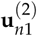 and the true genetic values **u**_*n*1_ is:

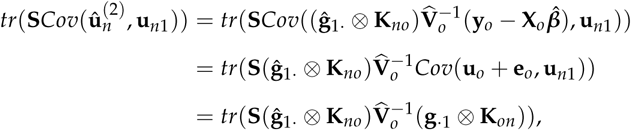

again assuming 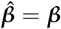 and now also *Cov* (**e**_*o*_, **u**_*n*1_) = **0**. The same result for the numerator of the expected correlation between 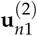 and the observed phenotypic values **y**_*n*1_ is:

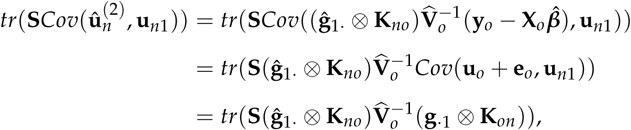

where we additionally assume *Cov*(**u**_*o*_, **e**_*n*1_) = **0** and *Cov* (**e**_*o*_, **e**_*n*1_) = **0**. Therefore, the numerators are the same, and 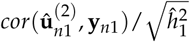 is a consistent estimator for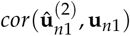.

### CV2-style multi-trait predictions

The numerator of the expected correlation between 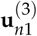 and the true genetic values **u**_*nl*_ is:

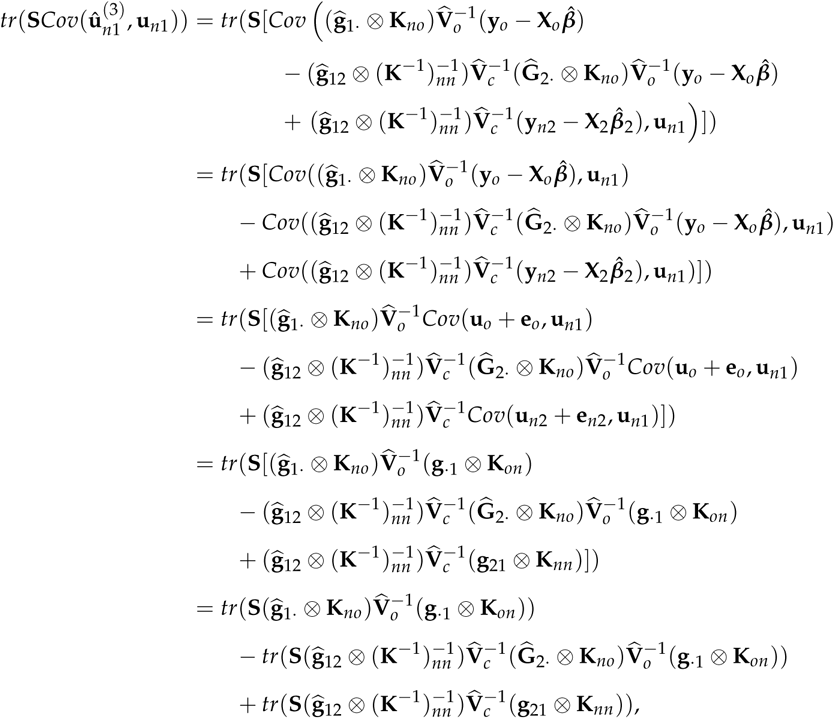

again assuming 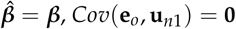, and *Cav(****e***_*n*2_, **u**_*n*1_) = **0**. From this, we can see the potential benefit of the CV2-style method:

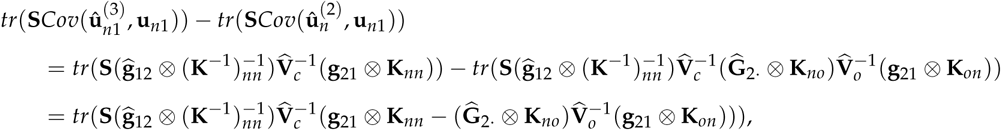

which is generally (but maybe not necessarily) positive. This means that 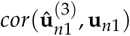 is generally greater than 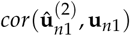.

The same result for the numerator of the expected correlation between 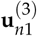 andtheobserved phenotypic values **y**_*n*1_ is:

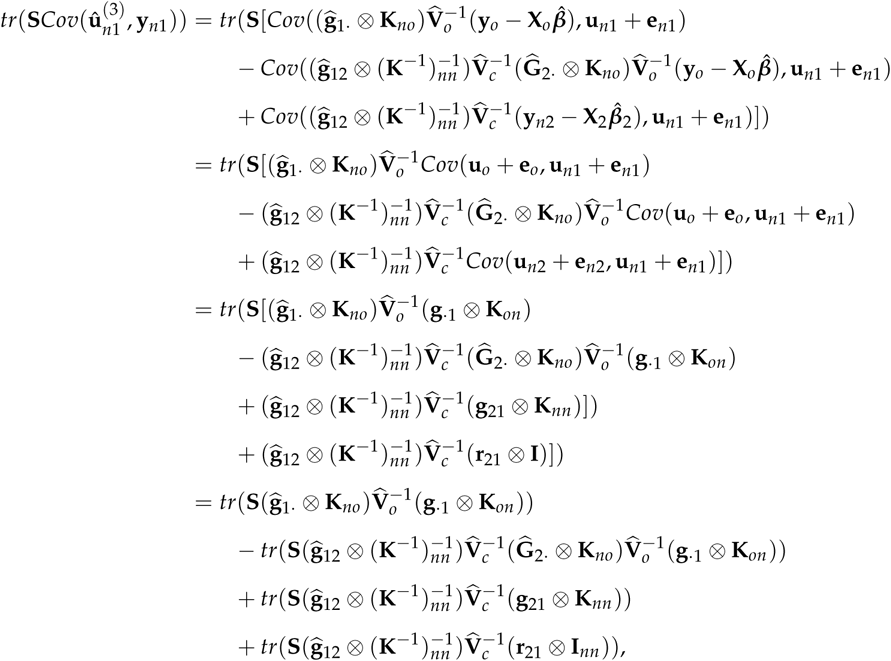

From this, we see that the numerator of the correlation 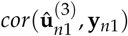 is not equal to that of 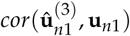:

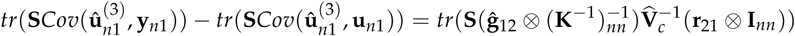

If *p* = 1, then **ĝ**_12_ and **r**_12_ are scalars and this excess covariance is approximately *n***ĝ**_12_**r**_12._

### CV2* approach

In our new CV2* cross-validation approach, we replace **y**_*n*l_ with **y**_*x*l_ -the phenotypes of a new set of individuals (x) that are relatives of the testing partition and were not part of the training partition. Let **K**_*xx*_ be the genetic relationships among these *n*_*x*_ individuals, and **K**_*xo*_ be their genetic relationships with the training partition. The numerator of the expected correlation 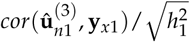 is:

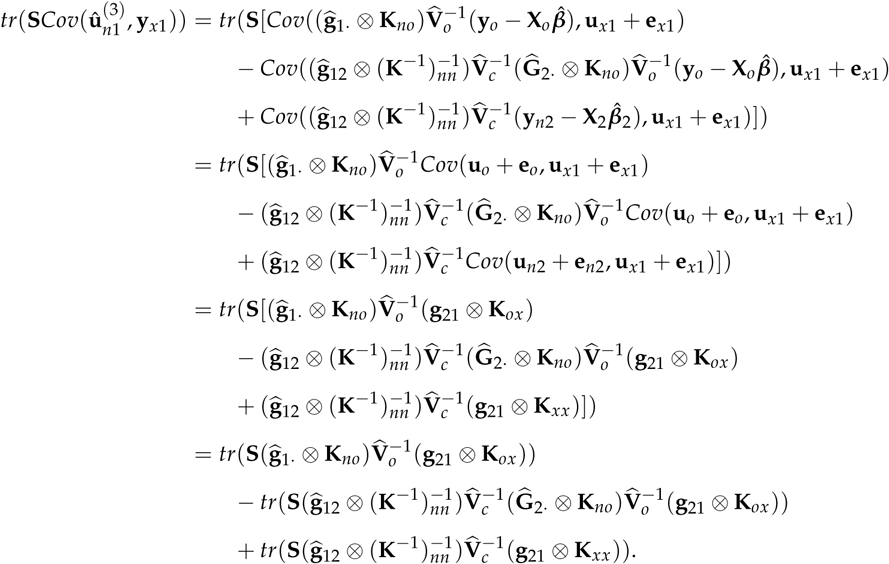

If these new individuals are clones of the original testing set, then **K**_*xx*_ = **K**_*nn*_, **K**_*ox*_ = **K**_*on*_ and 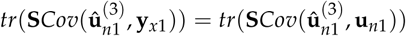 However, if clones are not available, then this equality will not hold.

Given these analytical results for the numerator of the expected correlations, we can estimate the correlation itself by calculating the expected variances of **û**_*n*1_ and **u**_*n*l_ or **y**_*n*l_. We do not go through these calculations as they follow directly from the calculations given above.

