## Supplementary figures and images for "Pitfalls and Remedies for Cross Validation with Multi-trait Genomic Prediction Methods"

### Supplemental Figure 1

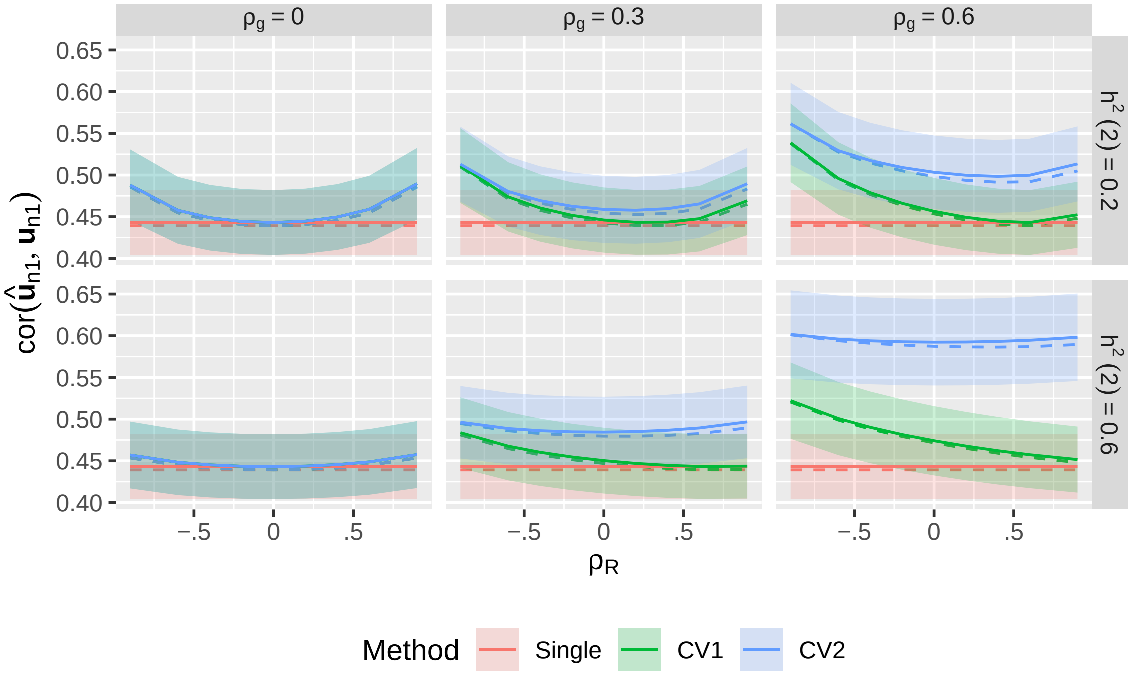

### Supplemental Figure 2

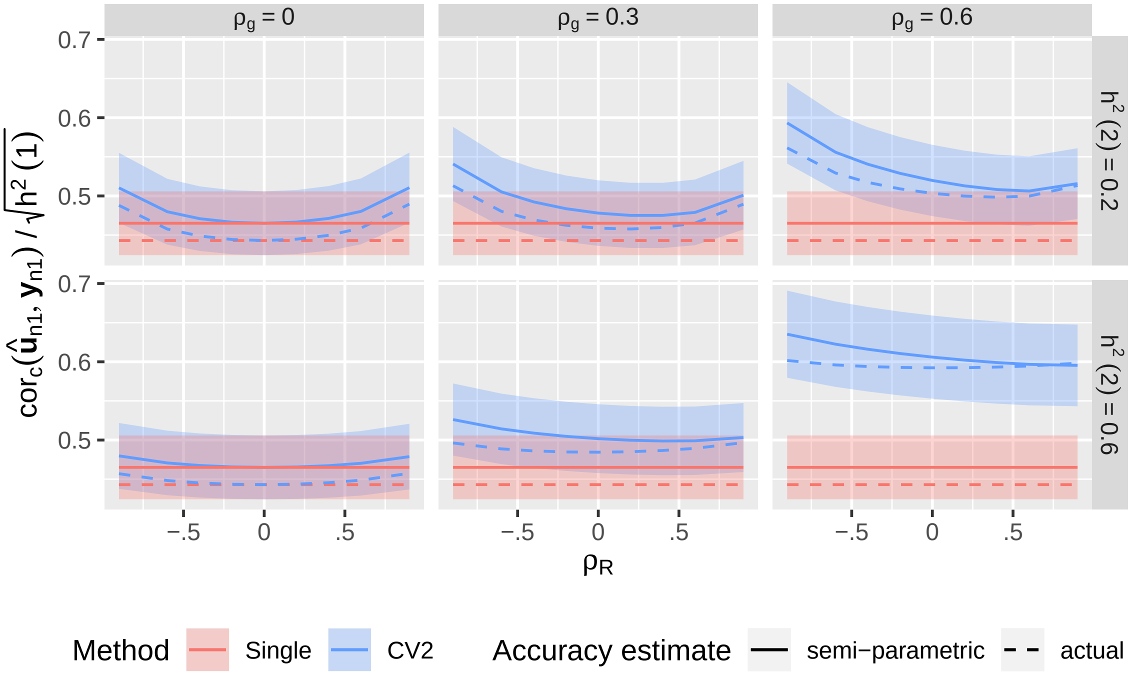
